# Extinction in Random Environments

**DOI:** 10.64898/2026.09.09.750454

**Authors:** Wenyun Zuo, Shripad Tuljapurkar

## Abstract

An important currency for individuals is their lifetime reproductive success (LRS), which is random simply due to demographic stochasticity. That randomness determines extinction probability. However, the distribution of LRS is also significantly affected by environmental variation. Previously we have shown (for a random environment that follows a Markov chain) how LRS is affected by an individual’s birth environment. But our previous analysis severs the temporal linkage between a parent’s birth environment and the environments into which its offspring are born. Here, we show how to compute the exact joint probability distribution of LRS for lineages spanning multiple environmental states (assuming a Markovian environment). From this joint distribution, we derive exact lineage extinction probabilities that fully incorporate demographic stochasticity, environmental frequency, and temporal autocorrelation. Applying our framework to Pacific Chinook salmon (semelparous with extreme early mortality) and European roe deer (iteroparous with delayed maturity), we demonstrate that initial birth states shape lineage fate. For salmon, the initial environment permanently separates trajectories; a poor birth state leads to near-certain extinction regardless of subsequent environmental shifts. For roe deer, we discover a counterintuitive dynamic where highly persistent poor conditions can rescue highly vulnerable individuals by expanding the right tail of reproduction. Furthermore, our exact calculations reveal that aggregated one-dimensional LRS distributions homogenize the reproductive landscape, overestimating extinction risks for lineages originating in poor environments and underestimating them for those in good environments. Accurately predicting evolutionary viability and the establishment of advantageous mutations requires preserving the environmental covariance. As global climate change amplifies environmental volatility, utilizing exact joint demographic models is critical for assessing true extinction risks.

## Introduction

Extinction is currently happening at an unprecedented rate, with many species disappearing due to human activities such as habitat destruction, climate change, pollution, over-fishing, and hunting (Tilman et al. 1994; Tilman and Lehman 1997; Nabi et al. 2018; Romero-Muñoz et al. 2021; Dulvy et al. 2021). In addition, the importance of internal and external stochasticity in environments is increasingly recognized as prevalent and impactful in ecology and evolution (Lenormand et al. 2009; Coulson 2021). Environmental conditions are changing and becoming increasingly variable and less predictable (IPCC 2023).

Even in a constant environment, extinction may occur because of bad luck! Any life history follows a sequence of intrinsically uncertain events, e.g., does an individual live or die, grow a little or a lot, have no or many offspring, have a few or many reproductive bouts? Recent work has shown that the accumulation of chance events in most life cycles leads to large variability in the Lifetime Reproductive Success (LRS) of individuals (Tuljapurkar et al. 2009; Caswell 2011; Steiner and Tuljapurkar 2012; van Daalen and Caswell 2015; Snyder and Ellner 2018; Tuljapurkar et al. 2020).

A family of population models with Branching Processes in Random Environments (BPRE) were used to study extinction probability (Vatutin et al. 2013). The general BPRE model was first introduced by Smith and Wilkinson (1969), and subsequently studied by Athreya and Karlin (1971b,a). Kozlov (1977), Vatutin and Dyakonova (1997), and Böinghoff et al. (2010) prove many asymptotic behaviors and limit theorems for BPRE. Haccou and Iwasa (1996) transform a population simulation into an efficient stochastic recurrence simulation by defining the extinction probability at time *t* based on the environment at that time and the extinction probability of the next generation at time *t* + 1. They find the variance and average extinction probability from the simulated stable extinction probability distribution. The average extinction probability does not distinct starting environment. However, all those models work with non-overlapping generations and discrete time steps in identically distributed random environments. Haccou and Vatutin (2003) extended the stochastic recurrence simulation to environmental autocorrelation but still only obtain the average extinction. Later, Pike et al. (2004) pointed out the methods of Haccou and Vatutin (2003) and Haccou and Iwasa (1996) could not calculate the generation-specific extinction probability for the species with overlapping generations.

To apply branching processes to life histories with overlapping generations, Tuljapurkar et al. (2020) developed a method to compute exactly the probability distribution of LRS, for any life history that, in discrete-time, is quantified by a stage-age-structured population model. Using the probability distribution of LRS, Tuljapurkar and Zuo (2022) made a general analysis of extinction probability in any structured population – even if the population grows on average, extinction may still occur with some probability. Extinction in their study means that starting with a single individual (with transition rates that would lead to long-run population increase *ω >* 1) but then that individual *and* all its descendants eventually go extinct (even when there is long-run population increase). They showed by examples that extinction probabilities can vary dramatically between life histories, depending on early survival and average fertility. They showed that the LRS distribution matters a great deal to the extinction probability of an advantageous mutant. The standard argument from Ewens (2012) to approximate the extinction probability using just the mean and variance of the LRS leads to wildly inaccurate results for some types of life history. For example, in semelparous species with massive maximum fertility like the Pacific Chinook Salmon, the massive variance in reproduction masks the inaccuracy of classical approximations, because extinction is close to certain. However, the traditional approximation under-estimates the actual chance of non-extinction by more than a factor of three.

To address the effect of fluctuating environmental conditions, due to factors such as climate change, Tuljapurkar et al. (2021) developed a method to compute the distribution of LRS in a random (Markovian, serially autocorrelated) environment. Using Roe deer (*Capreolus capreolus*) as a case study, they showed how the effect of an individual’s birth environment on LRS varies with the frequency of environments and their temporal autocorrelation. However, their method did not provide the joint probability distribution of offspring born in different environments, which is essential to calculate the extinction probability in varied environments.

Here, we begin by extending the method of Tuljapurkar et al. (2021) to provide the joint probability distribution of offspring born in different environments. We are then able to analyze the effect on extinction of both demographic stochasticity as well as environmental stochasticity (*e.g.*, distribution of environments, temporal autocorrelations). By avoiding simulation noise and approximation errors, our framework not only calculates exact extinction probabilities in autocorrelated random environments, but it also isolates the divergence in evolutionary fate between lineages starting in different environments. Using the Pacific Chinook salmon and the European roe deer as case studies, we demonstrate how environmental fluctuations interact with distinct birth states to shape the extinction and establishment probabilities of advantageous mutations, underscoring the absolute necessity of the joint LRS distribution.

### Finding the joint LRS

The Pacific Chinook salmon (*Oncorhynchus tshawytscha*) is a Pacific salmonid that breeds in the South Umpqua River in Oregon. The species is semelparous, meaning individuals reproduce only once in their lifetime and die immediately afterward. Individuals do not reproduce immediately; instead, they make their spawning run at age 3, 4, or 5 years. The population is censused annually, immediately before the spawning run. The projection matrix **A** has 7 stages, separating non-breeding ocean residents from breeding spawners to enforce the semelparous life history.

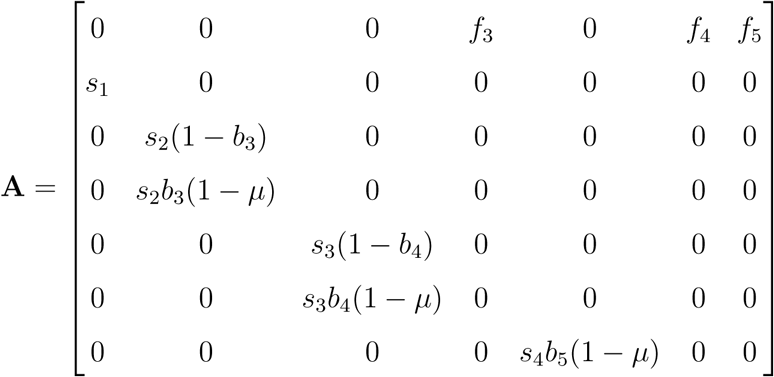

where *f_x_*is fecundity, the expected total number of eggs deposited by a successful spawner of age *x*, *s_x_* is ocean survival, the probability of surviving in non-breeding stages at age *x*, *b_x_* is breeding probability, the probability that an ocean resident of age *x* initiates the spawning run, *µ* is in-stream mortality, the additional mortality risk incurred during the upstream spawning migration (more details in appendix A1).

Decades of research have found that Pacific salmon populations have relatively stable adult ocean survival (*s_x>_*_1_) and fecundity (*f_x_*). The populations are primarily regulated by two environmental domains: ocean conditions during their first year, and river conditions during their spawning run. Ocean survival is dictated by the Pacific Decadal Oscillation (PDO) and El Niño Southern Oscillation (ENSO) (Mantua et al. 1997). Cool coastal waters and strong upwelling bring nutrient-rich waters to the surface, driving zooplankton blooms. This leads to rapid early growth and high first-year survival. El Niño events, and weak upwelling lead to nutrient depletion, smaller prey, and increased predation, causing massive early mortality (Michel et al. 2015). Freshwater survival is highly sensitive to river flow and temperatures (Crozier et al. 2008). Cool river temperatures and adequate flow rates allow for smooth passage to spawning grounds with low metabolic stress, which eases in-stream pre-spawning mortality.

We demonstrate our model by simply assuming that we have good or bad environments (*E_i_*, where *i* = 1 for good environment and *i* = 2 for bad) that affect *s*_1_(*E_i_*) (early ocean survival) and *µ*(*E_i_*) (in-stream mortality). There is a probability *ε_i_* that the environment is in state *i*, for an environmental autocorrelation *ψ*. Therefore we have an enviironment-dependent matrix population model (MPM) with matix **A**,

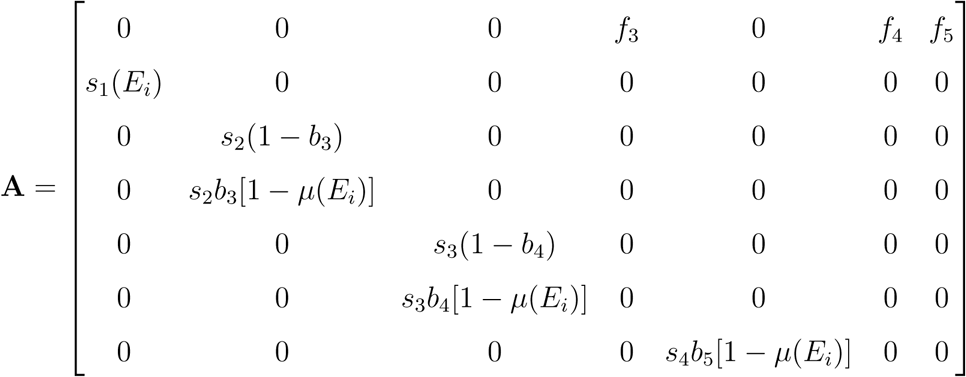

We want the exact joint probability, *P* (*k*_1_*, k*_2_), that an individual produces exactly *k*_1_ offspring in Environment 1 and *k*_2_ offspring in Environment 2 over their lifetime. Following the fundamental logic of Fast Fourier Transform (FFT) in the method derived by Tuljapurkar et al. (2020), the joint probability generating function *H*(*z*_1_*, z*_2_) for this individual’s entire lifetime looks like:

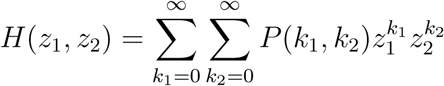

Let’s replace *z*_1_ and *z*_2_ with *θ*_1_ and*θ*_2_,

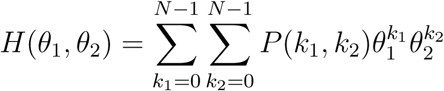

Now set

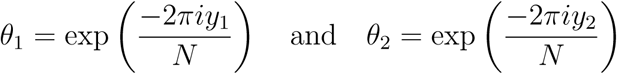

with two frequency indices, *y*_1_ and *y*_2_, each running from 0 to *N* − 1, where *N* is the maximum lifetime reproductive success, and turn our generating function into a two-deimensional (2D) discrete characteristic function

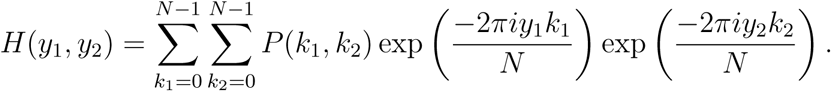

the exact value of the generating function *H*(*θ*_1_*, θ*_2_) evaluated at all frequencies. Then, we apply the 2D Inverse Fast Fourier Transform (IFFT) to get *P* (*k*_1_*, k*_2_).

### Onward to extinction probability in random environments

We can extend our previous Tuljapurkar and Zuo (2022) to two random environments. An offspring born in Environment 1, and all its descendants, goes extinct with probability *q*_1_, and one born in Environment 2 goes extinct with probability *q*_2_. Then we have a pair of nonlinear equation using the joint generating function at those probabilities:

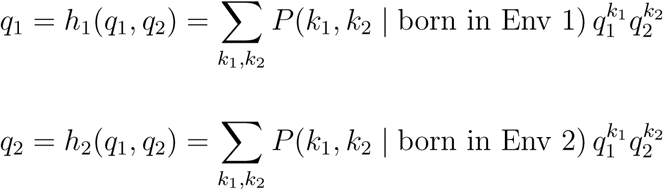

Since we can compute the 2D joint probability distributions for the offspring, we can calculate the exact extinction probability for an individual born in a specific environment.

### Case study with Pacific Chinook salmon

To parameterize the case study, we established environment-specific bounds for first-year survival and pre-spawning in-stream mortality. We defined first-year survival as *s*_1_(*E*_1_) = 0.003 for the good environment and *s*_1_(*E*_2_) = 0.0004 for the bad environment, derived from empirical smolt-to-adult return (SAR) data spanning two decades at the Coleman National Fish Hatchery (Adase et al. 2026) (details in Appendix A3). For pre-spawning in-stream mortality, we anchored our parameters around the commonly used baseline of *µ* = 0.1 (Crozier et al. 2008; Tuljapurkar and Zuo 2022). Informed by the official 2019–2020 California Department of Fish and Wildlife (CDFW) American River survey (Kelly and Phillips 2020), we set the mortality rates to *µ*(*E*_1_) = 0.05 for the good environment and *µ*(*E*_2_) = 0.25 for the bad environment (details in Appendix A4). The rest parameters in the MPM were kept identical to those in Tuljapurkar and Zuo (2022) (see Appendix A1).

For Pacific salmon, the first-year survival bottleneck exerts such disproportionate leverage on LRS that the initial environment permanently separates the lineage’s extinction trajectory. As illustrated in Figure 1, lineages originating in a Poor environment (*E*_2_, red crosses) face a near-certain extinction probability (*q >* 0.9998) that is insensitive to both the long-term frequency of good environments (*ε*_1_) and environmental autocorrelation (*ψ*). Because the baseline early mortality in the Poor environment is so severe (*s*_1_ = 0.0004), the initial cohort is effectively decimated before it can benefit from any subsequent shift to favorable conditions. Consequently, high autocorrelation only slightly worsen an already terminal trajectory. In contrast, lineages originating in a Good environment (*E*_1_, blue circles) successfully bypass this extreme initial bottleneck, rendering their ultimate viability highly dependent on the future environmental sequence. For these lineages, the extinction probability drops significantly as the long-term availability of favorable conditions (*π*_1_) increases. Furthermore, positive environmental autocorrelation (*ψ >* 0) substantially depresses *q* toward the theoretical minimum (fixed Good environment). The magnitude of the salmon’s initial mortality ensures that the birth state leaves an indelible mark on the lineage’s fate, even in an independent and identically distributed (i.i.d.) environment. Finally, it is critical to note the scale of the extinction probability (the y-axis): even under the most optimal stochastic scenarios (high *π*_1_, high *ψ*, born in *E*_1_), the ultimate extinction probability remains above 0.998, underscoring the extreme vulnerability inherent to this life-history strategy, which is due to extreme high initial mortality.

**Figure 1:**
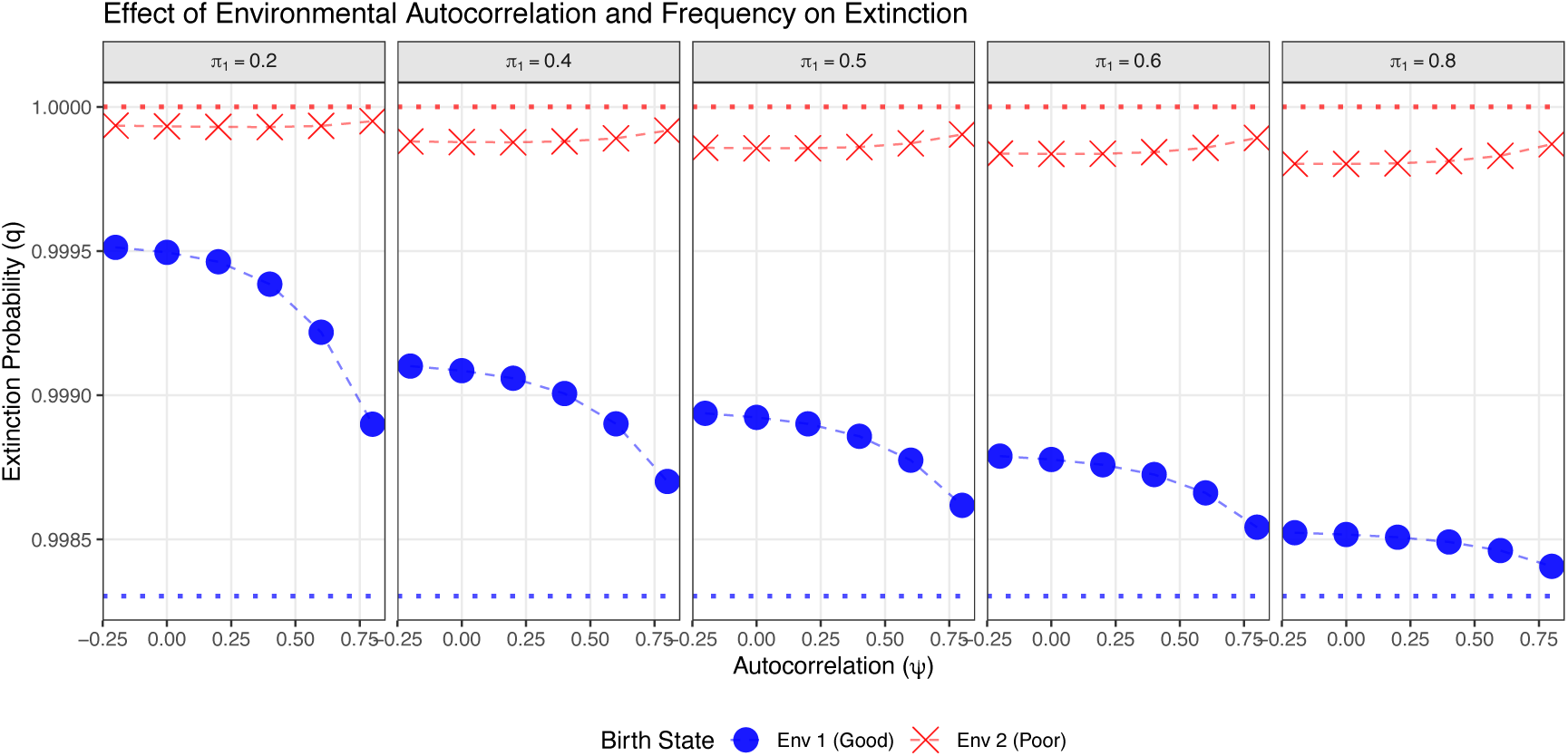
The interacting effects of environmental autocorrelation (*ψ*) and the frequency of favorable conditions (*π*_1_) on the extinction probability (*q*) of Pacific salmon lineages. Panels (left to right) represent an increasing long-term frequency of the Good environment (*π*_1_ ranging from 0.2 to 0.8). The x-axis denotes environmental autocorrelation (*ψ*), ranging from negative (alternating) to positive (persistent) environments. Data series indicate the environmental state at the lineage’s origin: blue circles represent a birth in the Good environment (*E*_1_), and red crosses represent a birth in the Poor environment (*E*_2_). The upper red dotted line indicates absolute extinction (*q* = 1), while the lower blue dotted line represents the theoretical minimum extinction probability (*q* = 0.9983) derived from a perpetually Good environment. The extreme first-year mortality bottleneck in the Poor environment causes the *E*_2_ trajectory to remain nearly flat and near *q* = 1 across all parameter space, while lineages starting in *E*_1_ are highly sensitive to both environmental persistence and long-term frequency.

### Case study with Roe Deer

We use the European roe deer (*Capreolus capreolus*) as another case study to demonstrate the stochastic environments effects and the birth state (body size) effects on lineage extinction probability (*q*). Tuljapurkar et al. (2021) used roe deer to demonstrate how random environmental fluctuations and distinct birth states shape the distribution of LRS. In the roe deer system, environmental states are defined by the timing of spring onset: “normal” springs represent Good environments (*E*_1_), whereas “early” springs act as Poor environments (*E*_2_). Empirical monitoring has demonstrated that during early spring years, the survival rates of young females (ages 1 to 7) are depressed to 90% of the baseline rates observed during normal springs. The corresponding integral projection model (IPM) captures high-resolution trait dynamics, tracking transitions across 12 discrete age classes and 200 body size classes. Continue with the life history used by Tuljapurkar et al. (2021), we calculated the joint probability of LRS and the lineage extinction probability.

Figure 2 is an example joint LRS distribution of a birth state at class 75 with the equal frequency of Good and Poor environments (*π*_1_ = 0.5) and a high environmental persistence (*ψ* = 0.8). The joint LRS distribution in this case is shaped by the initial birth environment. For individuals born into a Good environment, high autocorrelation ensures that their critical early survival and peak reproductive years likely align with continued favorable conditions. Consequently, their lifetime reproduction is heavily skewed toward producing offspring in the Good environment, expanding the probability mass along the *k_good_* axis and yielding a robust marginal probability of success. In the contrast, individuals born into a Poor environment face a compounding demographic penalty. Not only do they suffer depressed early survival – evidenced by the probability spike at the origin indicating reproductive failure – but those that do survive are trapped in a persistent sequence of poor years. As a result, their overall reproductive output is truncated, and the offspring they do manage to produce are predominantly born into the Poor environment, skewing the joint probability density vertically along the *k_bad_* axis. Compared to larger initial size classes, size class 25 exhibits prominent probability spikes at zero in the marginal plots (Figure A.2), indicating a much higher baseline risk of total reproductive failure. Despite this elevated early mortality, the high environmental persistence still visibly skews the successful reproductive mass toward the individual’s respective birth environment.

**Figure 2:**
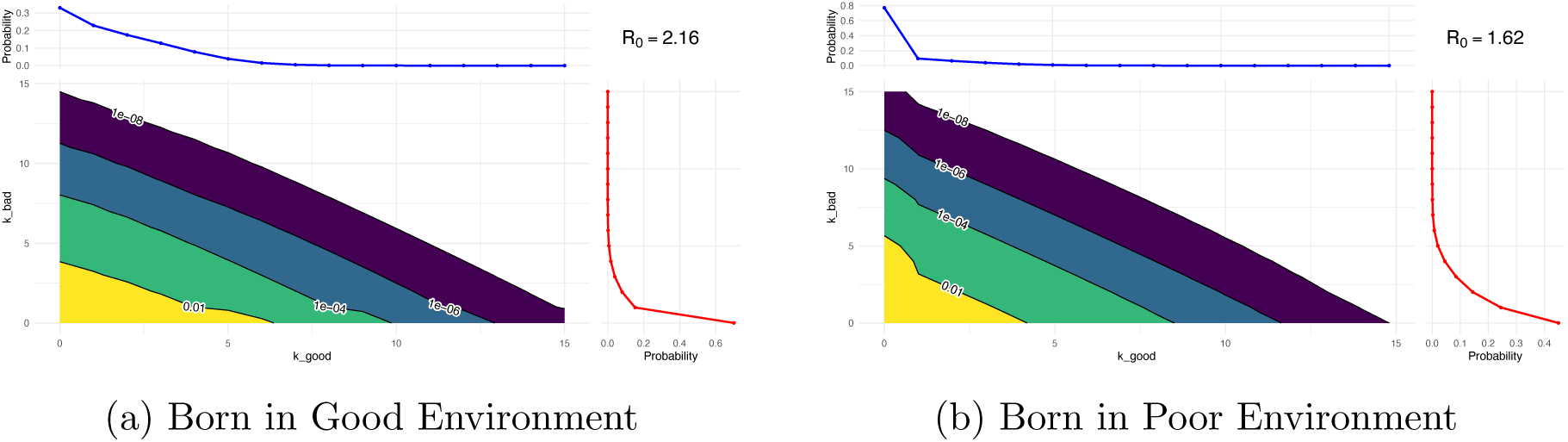
Joint Lifetime Reproductive Success (LRS) distributions for European roe deer originating in a Good environment (left) and a Poor environment (right). The central contour plots display the joint probability of an individual producing a specific combination of offspring in the Good environment (*k_good_*, x-axis) and the Bad environment (*k_bad_*, y-axis), with labeled contour lines denoting constant joint probability thresholds (ranging from 0.01 down to 10*^↑^*^8^). The attached marginal plots represent the probability densities of total offspring produced exclusively in the Good environment (top, blue line) and the Poor environment (right, red line). Both panels simulate a cohort starting at size class 75 under an equal long-term environmental frequency (*π*_1_ = 0.5) and high environmental persistence (*ψ* = 0.8).

The initial phenotypic state of an individual dictates its baseline extinction risk and alters how its lineage interacts with environmental stochasticity. To illustrate this, we compared the exact extinction probabilities for lineages originating from a robust initial size (size class 75, top panel) against those originating from a vulnerable size (size class 25, bottom panel). For larger newborns originating in a Poor environment, the lineage begins with a demographic buffer. Their baseline extinction risk is relatively low, but highly persistent Poor environments (*ψ >* 0) act as a demographic trap. Prolonged exposure to depressed survival conditions compounds mortality penalties, driving extinction trajectories upward. In contrast, lineages originating from smaller individuals in a Poor environment face a severe initial mortality bottleneck, shifting their baseline extinction risk upward to near certainty. However, an examination of the exact joint LRS matrices reveals a counterintuitive demographic tug-of-war for these vulnerable individuals. For a size 25 newborn originating in a Poor environment (at *π*_1_ = 0.4), high environmental persistence (*ψ* = 0.8) acts as a severe juvenile trap, increasing the probability of absolute reproductive failure (*P* (0, 0)) from 54.3% to 56.9% compared to a random fluctuating environment (*ψ*metic mean of the LRS (*R*_0_) down from 0.96 to 0.89. Despite this higher baseline mortality and lower average reproduction, the exact extinction probability of the highly autocorrelated regime is pulled below that of the fluctuating regime. This paradox is resolved by examining the extreme right tail of the joint LRS distribution, where the exact extinction probability (*q*) is highly sensitive to the environmental quality of “jackpot” reproductive events. As shown in Table 1, the persistent environment alters the probability of producing large, pure cohorts. Crucially, this expansion is asymmetric (Figure A.3). While the probability of producing 8 Poor-born offspring (*P* (0, 8)) increases 20-fold under high autocorrelation, the probability of producing 8 Good-born offspring (*P* (8, 0)) explodes by a factor of 68. The highly persistent environment guarantees that the tiny fraction of individuals surviving the initial Poor block are rewarded with a prolonged Good conditions, allowing them to produce concentrated streaks of high-quality offspring. Because Good-born offspring possess a higher probability of lineage establishment (*q*_1_ *< q*_2_), this asymmetric expansion of the right tail toward Good-born offspring anchors the generating function, overpowering the higher baseline juvenile mortality and rescuing the lineage trajectory.

**Table 1:** Reproductive Outcome Probabilities, *P* (*k*_good_*, k*_bad_) (sampled, full details in Appendix Figures A.2 and A.3), and mean LRS, *R*_0_, at Different *ψ* Levels on specific reproductive outcomes for a highly vulnerable European roe deer (size class 25) originating in a Poor environment (*π*_1_ = 0.4).

| | $\psi = 0$ | $\psi = 0.8$ | $P_{\psi=0.8}/P_{\psi=0}$ |
| --- | --- | --- | --- |
| $P(0, 0)$ | 0.543 | 0.569 | 1.05 |
| $P(0, 7)$ | $3.96 \times 10^{-5}$ | $5.18 \times 10^{-4}$ | 13 |
| $P(7, 0)$ | $3.34 \times 10^{-6}$ | $1.11 \times 10^{-4}$ | 33 |
| $P(0, 8)$ | $6.50 \times 10^{-6}$ | $1.28 \times 10^{-4}$ | 20 |
| $P(8, 0)$ | $3.69 \times 10^{-7}$ | $2.51 \times 10^{-5}$ | 68 |
| $P(4, 4)$ | $1.09 \times 10^{-4}$ | $3.71 \times 10^{-5}$ | 0.34 |
| $R_0$ | 0.96 | 0.89 | |

### Exact extinction probability and classical estimations

Classical demographic theory relies on the mean and variance of LRS to estimate lineage extinction probability, these approximations often fail to capture the true extinction dynamics (Tuljapurkar and Zuo 2022). To evaluate the limitations of traditional extinction risk models, we contrasted our exact extinction probabilities calculated from joint LRS against classical estimates derived from aggregated one-dimensional LRS statistics, mean and variance. As illustrated in Figure 4, the classical estimation method (purple markers), which relies solely on the mean and variance of the aggregated LRS, fails to capture the effects of structured, autocorrelated environments. While the exact extinction probabilities (blue circles for *E*_1_, red crosses for *E*_2_) exhibit a sensitivity to environmental persistence (*ψ*), diverging as regimes become more positively autocorrelated, the classical estimates remain flat across the entire *ψ* gradient. By collapsing the complex bivariate distribution of LRS into a single dimension and using only the mean and variance, the classical method homogenizes the temporal sequence of the environment. It smooths over the critical disadvantages of consecutive poor years and the advantages of consecutive favorable years that govern a lineage’s extinction trajectory.

**Figure 3:**
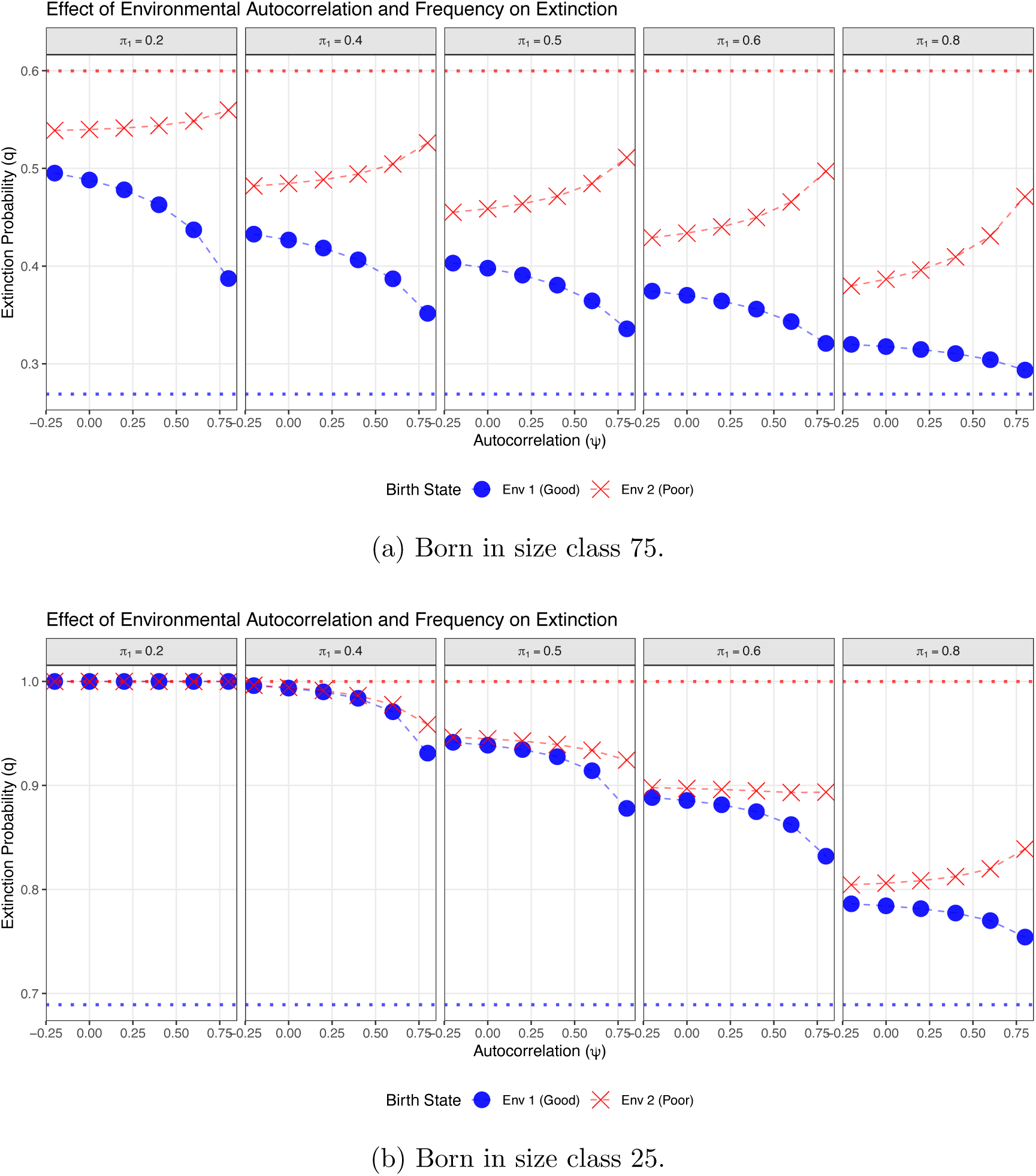
The interacting effects of initial size class and environmental autocorrelation (*ψ*) on exact lineage extinction probabilities for European roe deer. The top panel illustrates exact extinction trajectories for lineages originating as robust newborns (size class 75). The bottom panel displays trajectories for lineages starting as vulnerable newborns (size class 25). Columns represent an increasing long-term frequency of the Good environment (*π*_1_ ranging from 0.2 to 0.8). The x-axis denotes environmental autocorrelation (*ψ*). Blue circles (*E*_1_) represent lineages born into a Good environment, while red crosses (*E*_2_) represent those born into a Poor environment. The dotted lines represent theoretical extinction bounds derived from fixed environments. Note the largely different y-axis scales (*q*) between the panels, reflecting the severe baseline mortality penalty of a smaller initial size.

**Figure 4:**
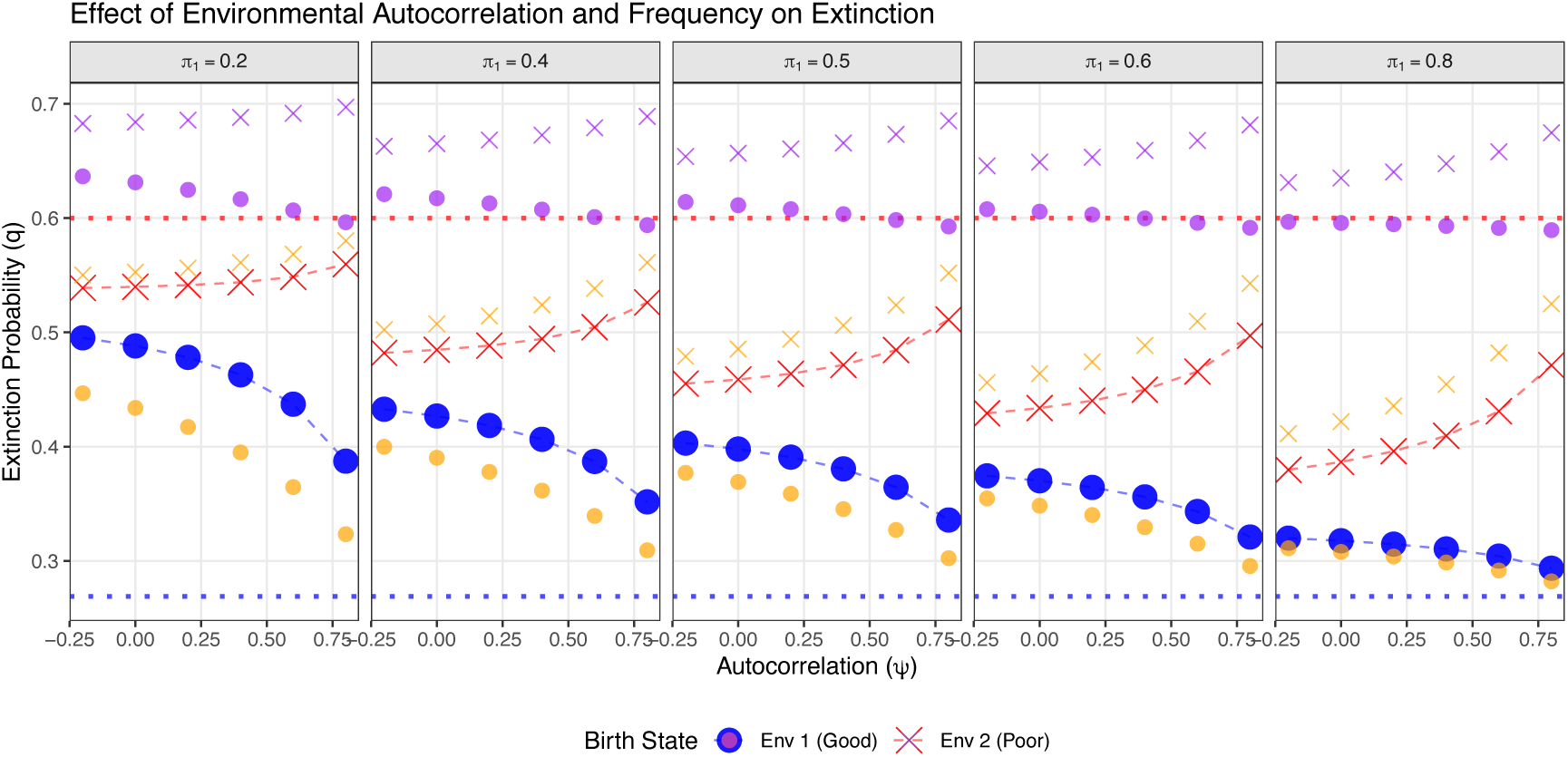
Comparison of exact extinction probabilities versus classical variance-based estimates for European roe deer lineages across varying environments. Roe deer born at the same birth size class 75. Panels (left to right) represent an increasing long-term frequency of the Good environment (*π*_1_ ranging from 0.2 to 0.8). The x-axis denotes environmental autocorrelation (*ψ*). Data point shapes indicate the birth environment: filled circles represent a birth in the Good environment (*E*_1_), and crosses represent a birth in the Poor environment (*E*_2_). Colors denote the calculation method: Blue (circles) and Red (crosses) represent the exact extinction probabilities derived from the joint LRS distribution. Purple markers represent the estimated extinction probability calculated via classical approximations utilizing the mean and variance of the aggregated LRS. Orange markers indicate the extinction probability calculated from the aggregated LRS. The upper red dotted line theoretical maximum extinction (*q* = 0.6) derived from a fixed Poor environment, while the lower blue dotted line represents the theoretical minimum extinction probability (*q* = 0.27) derived from a fixed Good environment.

To further evaluate the necessity of the joint LRS distribution, we compared our exact extinction probabilities against estimates derived from the aggregated one-dimensional (1D) LRS, utilizing the branching process methods established by Tuljapurkar and Zuo (2022). As illustrated in Figure 4, the aggregated 1D LRS estimation method (orange markers) represents a significant improvement over classical mean-variance approximations; it successfully captures the general shape of the trajectories driven by environmental autocorrelation (*ψ*) and the frequency of favorable conditions (*π*_1_). However, despite capturing the overall trends, collapsing the bivariate LRS distribution into a single dimension distorts the true magnitude of the extinction risk. Specifically, the aggregated 1D method overestimates the extinction risk for lineages originating in Poor environments (orange crosses higher than red crosses) while simultaneously underestimating the risk for lineages originating in Good environments (orange circles lower than blue circles). This distortion occurs because the 1D aggregation breaks the explicit temporal linkage between a parent’s current state and the specific environment into which its offspring are born. In the exact joint calculation, demographic pathways are preserved: a roe deer born in a Poor environment that manages to survive to maturity has a probability of producing offspring during a favorable environmental phase, buffering its lineage against failure. By blending these state-dependent pathways into a single aggregated distribution, the 1D method fails to capture this buffering. It assumes a homogenized reproductive landscape, causing it to exaggerate the penalties of the Poor environment and overstate the safety of the Good environment. This discrepancy clearly demonstrates that accurately predicting lineage viability requires not just the full distribution of LRS, but the explicit joint distribution that preserves environmental covariance.

Finally, by translating environmental autocorrelation into continuous temporal scales (see Appendix 8), our work captures the onset of demographic resonance, confirming that environmental impacts on extinction risk are amplified when environmental characteristic time aligns with a population’s generation time.

## Discussion

Our results demonstrate that environmental stochasticity dictates not just the immediate survival of a cohort, but the ultimate viability of the lineage across generations. Lineage survival cannot be accurately predicted using only average survival metrics or baseline mortality. Our exact calculations reveal that extinction probabilities are shaped by the interaction between the initial birth state, the temporal sequence of environmental fluctuations, and the extreme right tail of lifetime reproductive success. For iteroparous species with delayed reproduction like the European roe deer, the structural impact of environmental autocorrelation is modulated by the initial phenotypic state of the individual. For robust individuals starting increasingly persistent poor environments driving extinction trajectories upward. However, for vulnerable individuals, the exact calculation uncovers a counterintuitive dynamic: high environmental persistence surprisingly reduces extinction risk, even as it lowers the average reproduction (*R*_0_) and increases the reproduction failure rate (*P* (0, 0)) of the cohort. This rescue effect is driven entirely by an asymmetric variance in offspring quality. Vulnerable individuals born in highly persistent poor regimes experience a severe toll during its juvenile phase (increasing absolute lineage failure). However, highly persistent poor regimes in juvenile phase alter the reproductive landscape for the infinitesimal fraction of survivors. For those that reach peak reproductive size just as the environment shifts to a favorable state, high persistence (*ψ* = 0.8) guarantees a prolonged, uninterrupted window of optimal conditions. During this extended favorable block, these adults produce large cohorts of high-quality offspring (high lineage establishment cohorts). Conversely, rapidly fluctuating environments (*ψ* = 0) fail to provide this sustained favorable window; even if an individual survives to peak size, rapid environmental shifts constantly dilute their reproductive output, trapping the lineage in a cycle of mixed-quality offspring. Critically, our joint LRS framework reveals that blocky environments do not just expand the right tail of reproduction; they expand it asymmetrically, disproportionately inflating the probability of producing exclusive streaks of high-quality offspring. Because good-born offspring possess a higher baseline probability of lineage establishment, they essentially tilt the odds in favor of the lineage surviving.

Crucially, however, this rescue effect is heavily constrained by the individual’s initial phenotypic state and offers only a marginal relief from near-certain lineage failure. The asymmetric expansion of the right tail only overpowers the baseline mortality penalty for highly vulnerable individuals (e.g., size class 25) whose baseline extinction risk already borders on *q* = 1. For these individuals, the rare jackpot offers a slight but structural rescue, pulling the extinction probability down by a few percentage points. Yet, as the initial size of the individual increases and the baseline juvenile mortality bottleneck eases (e.g., reaching size class 50), this counterintuitive phenomenon rapidly fades out. For intermediate and robust individuals, the demographic buffer is already sufficient to prevent immediate lineage collapse. Consequently, the heavy demographic toll of enduring a highly persistent poor environment easily overwhelms the benefit of the right-tail jackpot, restoring the standard dynamic where high autocorrelation strictly increases extinction risk.

Central to accurately capturing these complex, state-dependent dynamics is the derivation of the joint LRS distribution. Classical demographic estimation homogenizes the temporal sequence of the environment, failing to capture the compounding structural impacts of autocorrelated regimes. While aggregated one-dimensional LRS methods successfully capture the general shape of extinction trajectories, collapsing the bivariate LRS distribution distorts the true magnitude of the extinction risk because it breaks the explicit temporal linkage between a parent’s demographic state and the specific environment of its offspring. The necessity of exact extinction probabilities extends directly to our understanding of evolutionary trajectories. Accurately predicting evolutionary viability and the establishment of advantageous mutations requires preserving this environmental covariance through the exact joint LRS distribution.

As global climate change amplifies environmental volatility, shifts in the timing and frequency of climatic fluctuations will become as consequential as changes in their magnitude. Our framework demonstrates that environmental impacts on extinction risk are amplified through demographic resonance, which occurs when the characteristic time of environmental cycles aligns with a population’s generation time (Appendix 8). Because even subtle phenotypic variations – such as the delayed maturation of a stunted roe deer – alter a lineage’s generation time, different phenotypes within the same population will become vulnerable to entirely different frequencies of environmental change. Predicting true evolutionary viability and the establishment of advantageous mutations in an increasingly stochastic world will therefore require moving beyond mean environmental trends to explicitly evaluate the temporal alignment between external climatic cycles and an organism’s intrinsic biological clock.

