## Appendix for "Extinction in Random Environments"

#### A1 MPM

The population is censused annually, immediately before the spawning run. The population state vector $\mathbf{n}(t)$ tracks individuals across 7 distinct stages, explicitly separating non-breeding ocean residents from breeding spawners to enforce the semelparous life history:

$$\mathbf{n}(t) = \begin{bmatrix} n_{1,O} \\ n_{2,O} \\ n_{3,O} \\ n_{3,S} \\ n_{4,O} \\ n_{4,S} \\ n_{5,S} \end{bmatrix}$$

Where $O$ denotes individuals remaining in the ocean and $S$ denotes individuals undertaking the spawning run.

The population is projected from year $t$ to $t+1$ using the equation $\mathbf{n}(t+1) = \mathbf{A}\mathbf{n}(t)$, where the projection matrix $\mathbf{A}$ isolates true fecundity in the top row and applies instream mortality explicitly to the spawner transitions:

$$\mathbf{A} = \begin{bmatrix} 0 & 0 & 0 & f_3 & 0 & f_4 & f_5 \\ s_1 & 0 & 0 & 0 & 0 & 0 & 0 \\ 0 & s_2(1-b_3) & 0 & 0 & 0 & 0 & 0 \\ 0 & s_2 b_3(1-\mu) & 0 & 0 & 0 & 0 & 0 \\ 0 & 0 & s_3(1-b_4) & 0 & 0 & 0 & 0 \\ 0 & 0 & s_3 b_4(1-\mu) & 0 & 0 & 0 & 0 \\ 0 & 0 & 0 & 0 & s_4 b_5(1-\mu) & 0 & 0 \end{bmatrix}$$

All demographic parameters are derived from the Pacific Chinook salmon (Oncorhynchus tshawytscha) life history.

Substituting the life history values into the symbolic matrix yields the exact nu-

Table 2: Demographic parameter definitions for the Pacific Chinook salmon (*Oncorhynchus tshawytscha*) life history model.

| Parameter | Value | Definition |
|---|---|---|
| $f_x$ | $f_3$: 3185 | **Raw Fecundity:** The expected total number of eggs deposited by a successful spawner of age $x$. |
| | $f_4$: 3940 | |
| | $f_5$: 4336 | |
| $s_x$ | $s_1$: 0.002267 | **Ocean Survival:** The probability of surviving non-breeding sources of mortality. $s_1$ encompasses all first-year mortality from deposited egg to an established Age 2 ocean resident. |
| | $s_{>1}$: 0.8 | |
| $b_x$ | $b_3$: 0.112 | **Breeding Probability:** The probability that an ocean resident of age $x$ initiates the spawning run. |
| | $b_4$: 0.532 | |
| | $b_5$: 1.0 | |
| $\mu$ | 0.1 | **In-Stream Mortality:** The additional mortality risk incurred during the upstream spawning migration. |

meric transitions.Notice that the columns corresponding to the Spawner stages ($n_{3,S}$, $n_{4,S}$, $n_{5,S}$) contain solely zeros beneath the top row, mathematically terminating those lineages after they deposit their raw fecundity (semelparity).

$$\mathbf{A} = \begin{bmatrix} 0 & 0 & 0 & 3185 & 0 & 3940 & 4336 \\ 0.002267 & 0 & 0 & 0 & 0 & 0 & 0 \\ 0 & 0.7104 & 0 & 0 & 0 & 0 & 0 \\ 0 & 0.08064 & 0 & 0 & 0 & 0 & 0 \\ 0 & 0 & 0.3744 & 0 & 0 & 0 & 0 \\ 0 & 0 & 0.38304 & 0 & 0 & 0 & 0 \\ 0 & 0 & 0 & 0 & 0.72 & 0 & 0 \end{bmatrix}$$

### A2 Diagonal matrix, $\tilde{\mathbf{W}}(y_i)$

Where the fully expanded 14-element diagonal is:

$$\tilde{\mathbf{W}}(y_i) = \text{diag}\begin{pmatrix} 1 \\ 1 \\ 1 \\ \tilde{w}_3(y_1) \\ 1 \\ \tilde{w}_4(y_1) \\ \tilde{w}_5(y_1) \\ 1 \\ 1 \\ 1 \\ \tilde{w}_3(y_2) \\ 1 \\ \tilde{w}_4(y_2) \\ \tilde{w}_5(y_2) \end{pmatrix}$$

### A3 The combined stage transition matrix, Q

The environments transition matrix

$$\mathbf{P} = \begin{bmatrix} P_{11} & P_{12} \\ P_{21} & P_{22} \end{bmatrix}$$

The combined stage transition matrix,

$$\mathbf{Q} = \begin{bmatrix} P_{11}\mathbf{U}_1 & P_{12}\mathbf{U}_2 \\ P_{21}\mathbf{U}_1 & P_{22}\mathbf{U}_2 \end{bmatrix}$$

Where the transition matrices ($\mathbf{U}_1$ in good environment and $\mathbf{U}_2$ in bad environment) are

$$\mathbf{U}_1 = \begin{bmatrix} 0 & 0 & 0 & 0 & 0 & 0 & 0 \\ 0.003 & 0 & 0 & 0 & 0 & 0 & 0 \\ 0 & 0.7104 & 0 & 0 & 0 & 0 & 0 \\ 0 & 0.08512 & 0 & 0 & 0 & 0 & 0 \\ 0 & 0 & 0.3744 & 0 & 0 & 0 & 0 \\ 0 & 0 & 0.40432 & 0 & 0 & 0 & 0 \\ 0 & 0 & 0 & 0 & 0.76 & 0 & 0 \end{bmatrix}$$

and

$$\mathbf{U}_2 = \begin{bmatrix} 0 & 0 & 0 & 0 & 0 & 0 & 0 \\ 0.0004 & 0 & 0 & 0 & 0 & 0 & 0 \\ 0 & 0.7104 & 0 & 0 & 0 & 0 & 0 \\ 0 & 0.0672 & 0 & 0 & 0 & 0 & 0 \\ 0 & 0 & 0.3744 & 0 & 0 & 0 & 0 \\ 0 & 0 & 0.3192 & 0 & 0 & 0 & 0 \\ 0 & 0 & 0 & 0 & 0.60 & 0 & 0 \end{bmatrix}.$$

### A4 Parameterization of Environmental Stochasticity

To evaluate the evolutionary consequences of environmental stochasticity on population viability, we adapted the deterministic Chinook salmon matrix population model from Tuljapurkar and Zuo (2022) into a two-state Markov chain representing "Good" (highly productive) and "Bad" (resource-limited) ocean regimes. We introduced environmental variance exclusively into the first-year marine survival parameter ($s_1$), as early ocean entry represents the primary demographic bottleneck driving interannual recruitment volatility. The variance envelope was bounded using empirical smolt-to-adult return (SAR) data spanning two decades from the Coleman National Fish Hatchery (Adase et al. 2026). For the "Bad" environmental state, which simulates severe climate stressors such as marine heatwaves and prolonged drought, we defined first-year survival as $s_1 = 0.0004$. This lower bound explicitly reflects the catastrophic 0.03% SARs recorded during the extreme 2007 and 2020 outmigration years, assuming a standard late-stage adult ocean survival probability of 0.80. Conversely, for the Good environmental state, representing optimal, cold-water coastal upwelling conditions, we established an upper bound of $s_1 = 0.003$. This value aligns with the maximum peak return rates observed within the same empirical dataset. By stretching the first-year survival parameter across this biologically realistic, order-of-magnitude spread, the model accurately replicates the sweepstakes reproductive dynamics inherent to Pacific salmon and ensures that the variance penalty on the long-term stochastic growth rate is robustly captured.

In the official 2019–2020 CDFW American River survey (Kelly and Phillips 2020), a total of 3,042 female carcasses were assessed. 679 of those females were "unspawned" ($> 70\%$ egg retention). So the established pre-spawning mortality rate for that year was 22%.

#### A5 Exact extinction probability and classical estimations in Salmon case

To evaluate the limitations of traditional extinction risk models, we contrasted our exact extinction probabilities calculated from joint LRS against classical estimates derived from aggregated one-dimensional LRS statistics, mean and variance. The main text has shown the Roe deer case. Here we provide the results for Salmon case. As illustrated in Figure A.1, the classical estimation method (purple markers), which relies solely on the mean and variance of the aggregated LRS, fails to capture the effects of structured, autocorrelated environments. While the exact extinction probabilities (blue circles for $E_1$, red crosses for $E_2$) exhibit a sensitivity to environmental persistence ($\psi$), diverging as regimes become more positively autocorrelated, the classical estimates remain flat across the entire $\psi$ gradient. By collapsing the complex bivariate distribution of LRS into a single dimension and using only the mean and variance, the classical method homogenizes the temporal sequence of the environment. It smooths over the critical disadvantages of consecutive poor years and the advantages of consecutive favorable years that govern a lineage's extinction trajectory.

#### A6 Example of roe deer birth state at class 25

To illustrate how demographic vulnerability interacts with environmental stochasticity, we examined the joint LRS for European roe deer originating at a smaller initial size (size class 25, Figure A.2). Because smaller individuals face higher pre-reproductive mortality, the dominant feature of these distributions is the massive probability spike at the origin ($k_{good} = 0$, $k_{bad} = 0$) across both birth states. For these smaller individuals, demographic stochasticity dominates the early life history, resulting in a high probability of total reproductive failure. However, for the fraction of the cohort that survives this initial, size-dependent mortality, the highly persistent environmental regime ($\psi = 0.8$) still enforces a strict divergence in lifetime trajectories. Individuals born in a Good environment that successfully reach maturity skew their reproductive output along the $k_{good}$ axis, benefiting from consecutive favorable years. Conversely, individu-

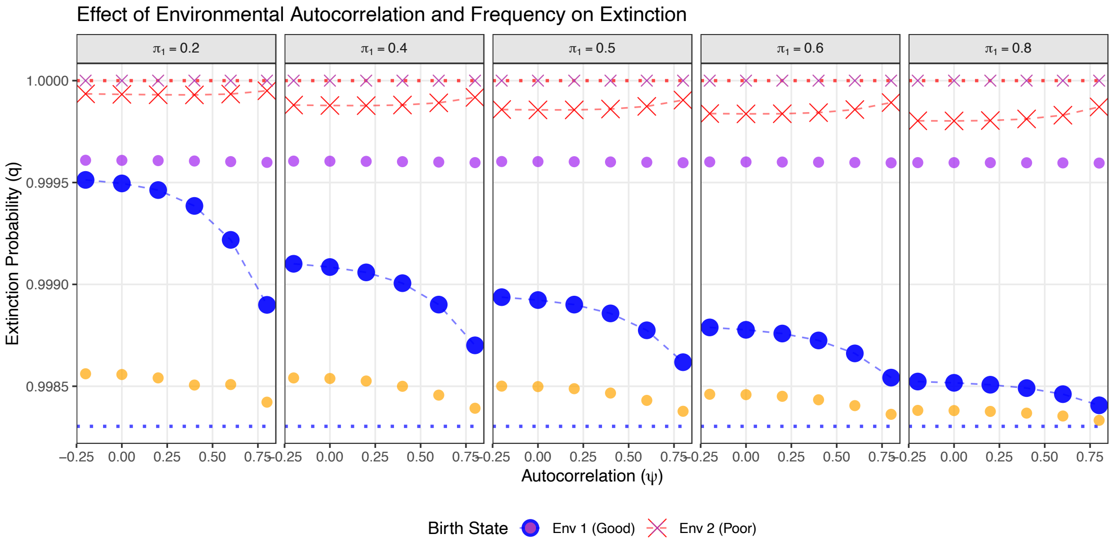


Figure A.1: Comparison of exact extinction probabilities versus classical variance-based estimates for Pacific Salmon lineages across varying environments. Panels (left to right) represent an increasing long-term frequency of the Good environment ($\pi_1$ ranging from 0.2 to 0.8). The x-axis denotes environmental autocorrelation ($\psi$). Data point shapes indicate the birth environment: filled circles represent a birth in the Good environment ($E_1$), and crosses represent a birth in the Poor environment ($E_2$). Colors denote the calculation method: Blue (circles) and Red (crosses) represent the exact extinction probabilities derived from the joint LRS distribution. Purple markers represent the estimated extinction probability calculated via classical approximations utilizing the mean and variance of the aggregated LRS. Orange markers indicate the baseline probability of reproductive failure ($P(LRS = 0)$). The upper red dotted line indicates absolute extinction ($q = 1$) derived from a fixed Poor environment, while the lower blue dotted line represents the theoretical minimum extinction probability ($q = 0.9983$) derived from a fixed Good environment.

als originating in a Poor environment suffer a compounded demographic penalty: their already elevated size-based mortality is heavily exacerbated by the persistent poor conditions. This dual penalty truncates their total lifetime output, compressing the contour lines tightly toward the origin and confining what little reproduction they do achieve predominantly to the Poor environmental state.

The persistent environment alters the probability of producing large, pure cohorts. This expansion is asymmetric (Figure A.3). The contour plot displays the relative multiplier effect ($P_{\psi=0.8}/P_{\psi=0}$) of environmental autocorrelation on specific reproductive outcomes for a highly vulnerable European roe deer (size class 25) originating in a Poor environment ($\pi_1 = 0.4$). The dark, unlabeled central region spanning the diagonal represents mixed-quality reproductive outcomes (e.g., $k_{good} \approx k_{bad}$) whose probabilities are suppressed (ratio $< 1$) by high autocorrelation. Conversely, the asymmetric expansion of the contours along the x-axis confirms the right-tail "jackpot" effect. High environmental persistence ($\psi = 0.8$) disproportionately inflates the probability of producing exclusive streaks of high-quality, Good-environment born offspring by over two orders of magnitude (the 100+ contour region) compared to a rapidly fluctuating environment ($\psi = 0$).

### A7 Exact extinction probability and the characteristic time

Autocorrelation $\rho$ gives the probability that at time $t$ the environment is the same as current is $\rho^t$. Therefore,

$$\rho^t = e^{-|ln\rho|t} = e^{-t/\tau_E},$$

which gives the environmental characteristic time

$$\tau_E = \frac{1}{|ln\rho|}.$$

When the characteristic time scale of the environment ($\tau_E$) aligns with the generation time ($T_c$), the demographic impact of the environment is amplified. This phenomenon has been referred as demographic resonance (or cohort resonance) (Worden et al. 2010).

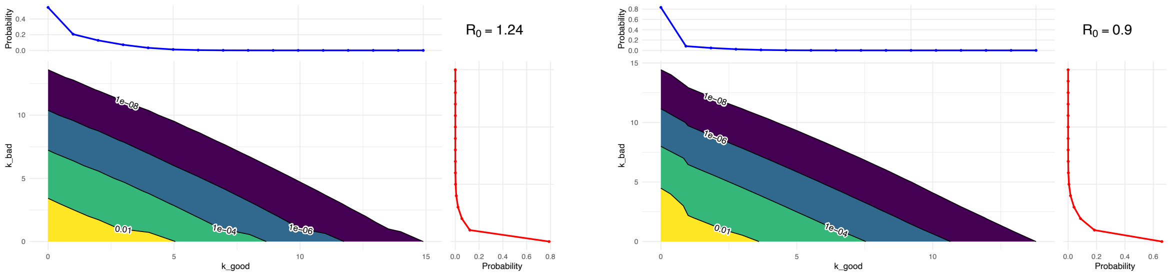


(a) Born in Good Environment (b) Born in Poor Environment

Figure A.2: Joint Lifetime Reproductive Success (LRS) distributions for European roe deer starting at a smaller initial size (size class 25) originating in a Good environment (left) and a Poor environment (right). The central contour plots display the joint probability of an individual producing a specific combination of offspring in the Good environment ($k_{good}$, x-axis) and the Bad environment ($k_{bad}$, y-axis), with labeled contour lines denoting constant joint probability thresholds. The attached marginal plots represent the probability densities of total offspring produced exclusively in the Good environment (top, blue line) and the Poor environment (right, red line). Both panels simulate a cohort starting at size class 25 under an equal long-term environmental frequency ($\pi_1 = 0.5$) and high environmental persistence ($\psi = 0.8$). Compared to larger initial size classes, size class 25 exhibits prominent probability spikes at zero in the marginal plots, indicating a much higher baseline risk of total reproductive failure. Despite this elevated early mortality, the high environmental persistence still visibly skews the successful reproductive mass toward the individual's respective birth environment.

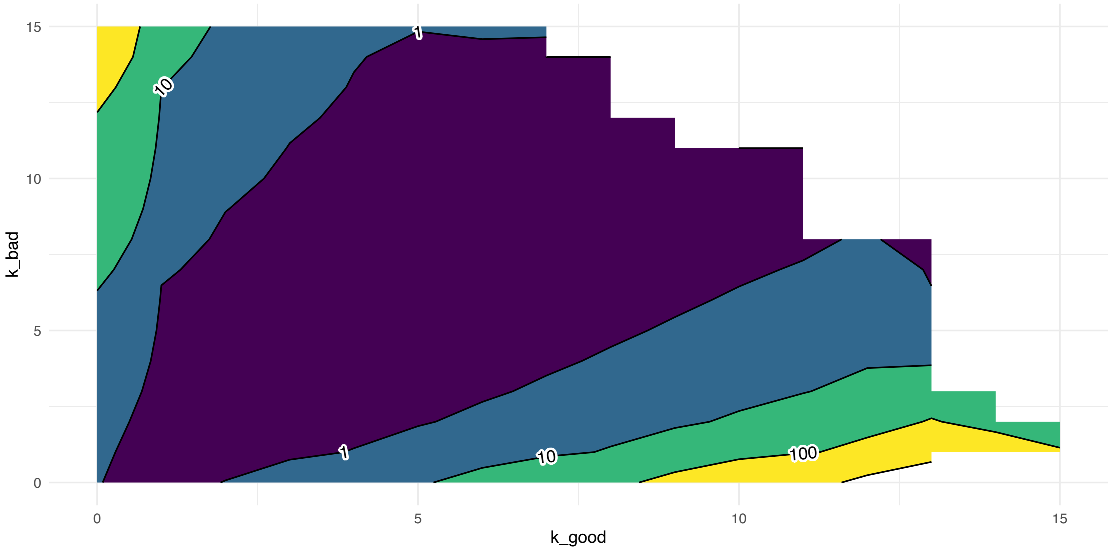


Figure A.3: Ratio of joint Lifetime Reproductive Success (LRS) probabilities between highly persistent and random fluctuating environments. The contour plot displays the relative multiplier effect ($P_{\psi=0.8}/P_{\psi=0}$) of environmental autocorrelation on specific reproductive outcomes for a highly vulnerable European roe deer (size class 25) originating in a Poor environment ($\pi_1 = 0.4$). The x-axis denotes offspring produced in the Good environment ($k_{good}$), and the y-axis denotes offspring produced in the Poor environment ($k_{bad}$). Contour lines designate boundaries of constant probability ratios (1, 10, 100 and 1000). The dark, unlabeled central region spanning the diagonal represents mixed-quality reproductive outcomes (e.g., $k_{good} \approx k_{bad}$) whose probabilities are heavily suppressed (ratio $< 1$) by high autocorrelation. Conversely, the asymmetric expansion of the contours along the x-axis confirms the right-tail "jackpot" effect. High environmental persistence ($\psi = 0.8$) inflates the probability of producing exclusive streaks of high-quality, Good-environment born offspring by over two orders of magnitude (the 100+ contour region) compared to a rapidly fluctuating environment ($\psi = 0$). Empty place in the figure represents zero probability in the joint LRS so the ratio is NA.

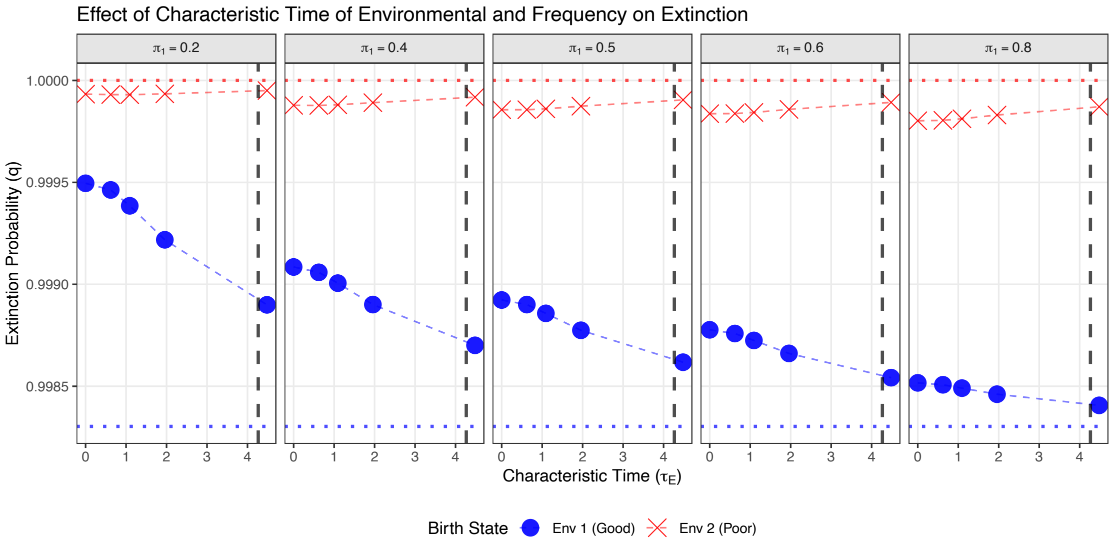


Figure A.4: The extinction probability ($q$) is plotted against the characteristic time of the environment ($\tau_E$) across a gradient of environmental frequencies ($\pi_1$, representing the proportion of time spent in the Good environment). Blue circles track the extinction probability for lineages originating in the Good environment (Env 1), while red crosses denote lineages originating in the Poor environment (Env 2). The vertical dashed line marks the generation time ($T_c \approx 4.2$ years) for the population. Under conditions of rapid environmental fluctuation ($\tau_E \ll T_c$), the outcomes for both starting states remain tightly coupled. However, as the characteristic time of the environment approaches the generation time ($\tau_E \approx T_c$), the divergence between the Good and Poor starting states maximizes. This expanding vertical gap demonstrates the onset of demographic resonance.

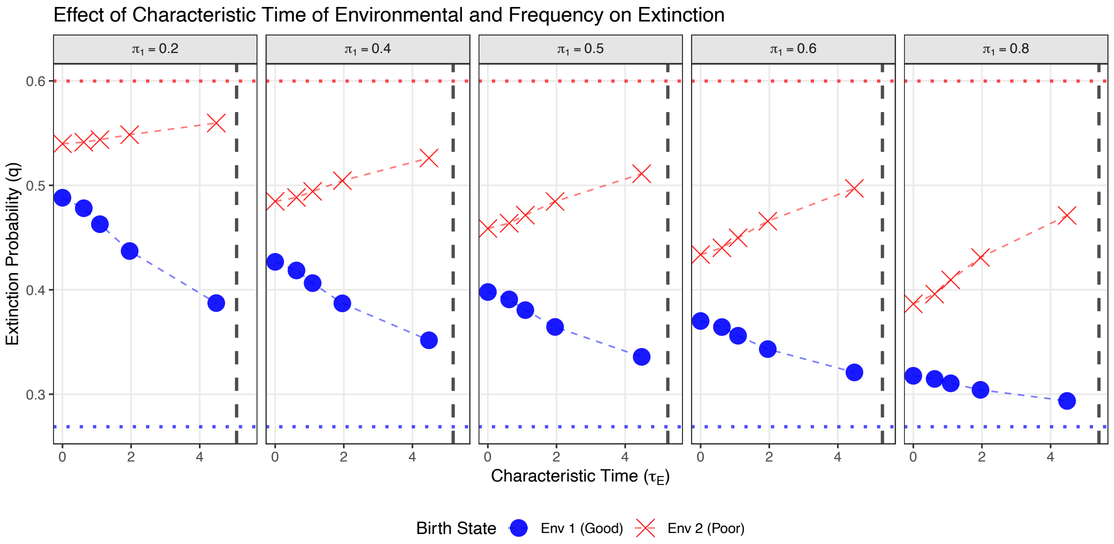


(a) Born in size class 75.

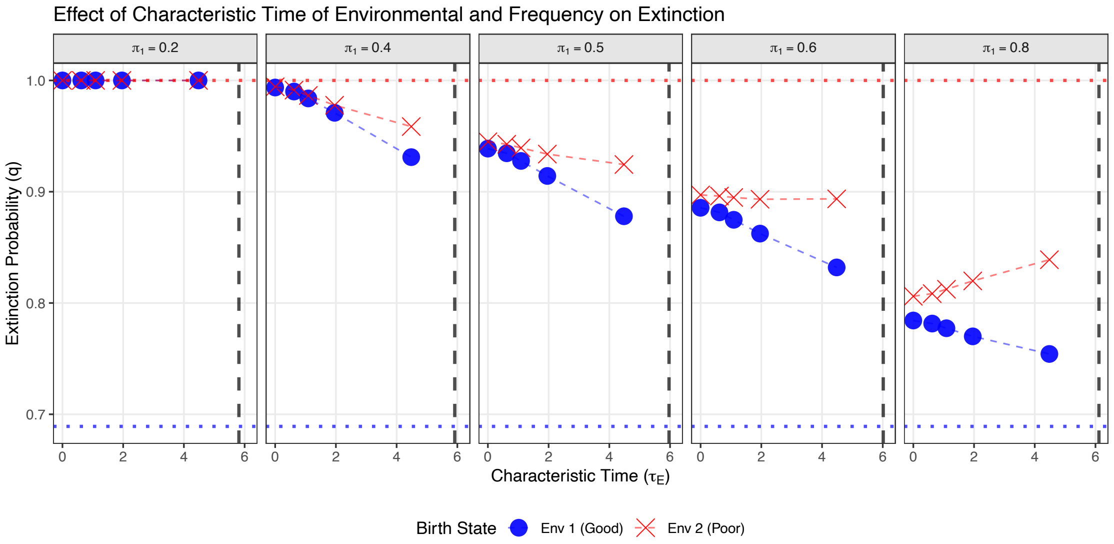


(b) Born in size class 25.

Figure A.5: Extinction probability ($q$) plotted against environmental characteristic time ($\tau_E$) across a gradient of Good environment frequencies ($\pi_1 \in [0.2, 0.8]$) for (a) robust size-class 75 newborns ($T_c \approx 4.8$ years) and (b) stunted size-class 25 newborns ($T_c \approx 5.9$ years). Blue circles indicate lineages originating in the Good environment (Env 1), while red crosses denote lineages originating in the Poor environment (Env 2). Vertical black dashed lines mark the generation time ($T_c$) calculated for each respective birth state. At rapid environmental fluctuations ($\tau_E \ll T_c$), initial birth environments are strongly buffered, resulting in tightly clustered extinction probabilities. As environmental persistence increases ($\tau_E \to T_c$), demographic resonance occurs: the vertical divergence between Good and Poor starting states expands to its maximum gap near $T_c$, demonstrating that environmental shock is heavily amplified when environmental cycles align with the population's reproductive pace. Comparing (a) and (b) reveals that robust founders compress the generation time by bypassing early growth bottlenecks, shifting the peak resonance zone to shorter environmental time scales while maintaining substantially lower overall extinction probabilities across all frequencies.
